# Moleculewise semi-grand canonical ensembles

**DOI:** 10.1101/2020.10.30.362178

**Authors:** M. Girard, T. Bereau

## Abstract

The plasma membrane is the interface between cells and exterior media. While its existence has been known for a long time, organization of its constituent lipids remain a challenge. Recently, we have proposed that lipid populations may be controlled by chemical potentials of different lipid species, resulting in semi-grand canonical thermodynamic ensembles. However, the currently available molecular dynamics software packages do not allow for molecule-based chemical potentials. Here, we propose a variation on existing algorithms that allow defining chemical potentials for molecules. Additionally, we allow coupling with collective variables and show that it can be used to dynamically create asymmetric membranes. We release an implementation of the algorithm for the HOOMD-Blue molecular dynamics engine.

**SIGNIFICANCE:** We demonstrate an algorithm that allows for simulations of molecules in the semi-grand canonical ensemble. It also allows coupling the chemical potential values to collective variable and create asymmetric membranes.

## INTRODUCTION

Membranes in eukaryote cell are mostly comprised of lipids, with particularly complex chemistry and organization. A typical mammal cell has hundreds of different lipids types in any of its membranes, distributed asymmetrically between both leaflets (1,2). The chemical nature of lipids — overall headgroup composition, acyl tail length, unsaturation — is maintained by the Lands’ cycle in the endoplasmic reticulum. The asymmetric distribution is maintained by type IV P-type ATPa_SE_ (P4-ATPA_SE_) proteins, also known as flippases, embedded in the membrane itself, which consume ATP in order to move lipids from one leaflet to the other. Given the length to which cells go to maintain their lipid composition, one can ask: why do cells require such a complex chemistry ? Computer simulations have proven excellent to garner insights into behavior of simple model membranes, and is moving towards realistic biological chemistry (3). For instance, the MARTINI model (4) has been used to model realistic simulations of plasma membranes (5). However, understanding the underlying fundamental reasons for membrane composition and asymmetry requires systematic variations of the myriad of potential compositions. Moreover, simulations involving asymmetric compositions must be done carefully as differential stress can exist in the membrane (6).

A related question is: how do membrane regulate their composition ? Giant plasma membrane vesicles—vesicles extracted from plasma membranes that retain composition— are known to possess a miscibility transition temperature just under cell growth temperature (7), which is in all appearance critical (8), clearly showing that lipid composition is responsive to environmental changes. Computer simulations are moving towards biologically relevant compositions (3); yet are still unable to correctly model regulation as it involves chemical reactions and lipid diffusion between membranes in cells. The problem appears enigmatic in experiments as well: no sophisticated sensing mechanism has been observed to precisely control the large amount of lipids types in membranes. We recently hypothesized that regulation of phospholipids in cells may be loose, and controlled by their chemical potential, while other components such as cholesterol may be tightly regulated (9). We named this configuration regulated ensembles, and it thermodynamically corresponds to mixtures of canonical and semi-grand canonical (SGC) ensembles — the thermodynamic ensemble where chemical potential differences between molecules is fixed. In simulations, some components can change their chemical nature over time, while their overall number is constrained. Subsequently, we have shown that this naturally self-regulates towards critical points, in a robust fashion (10).

Here, we present the software we employed to simulate lipids in SGC ensembles in (9). There already exists a highly parallel algorithm for SGC (11), available in multiple molecular dynamics packages, e.g. LAMMPS and openMM (12). However, it lacks two features for membrane simulations. First, it is unable to capture chemical potentials of molecules. Second, it does not allow coupling to collective variables. This is important for biological membranes as chemical potentials on the two leaflets are different. In order to resolve this, we extend the method to associate a chemical potential to any arbitrary combination of chemistry, charge state and collective variable. We release an implementation running on graphical processing units in the HOOMD-Blue molecular dynamics engine (13, 14). We use this implementation to simulate a lipid bilayer with an asymmetric composition. In order to do this, we postulate that P4-ATPA_SE_ induces a chemical potential difference that only depends on headgroup nature (phosphatidylcholine (PC) vs phosphatidyletholamine (PE)). This proxy allows us to dynamically create the asymmetry and relate the work done by P4-ATPA_SE_ on lipids to create the asymmetric profile.

## METHODS

The algorithm employed here makes use of a simulation domain checkerboard decomposition (see Fig 1A) in the same fashion as (11). The simulation box is decomposed into cells of minimal thickness *σ*, where *σ* is the largest interaction range in the system. Particles located at least two cells away from each other are therefore non-interacting. Every update step, the algorithm selects a set of non-interacting cells (depicted in blue in Fig 1A). Within this set of active cells, one particle is randomly selected and a swap is attempted, with acceptance determined by the usual Metropolis criterion exp(–*β*(Δ*U* – Δ*μ*)), where *β* = (*k_B_T*)^-1^, Δ*U* is the internal energy change and Δ*μ* the chemical potential difference between the two species.

**Figure 1:**
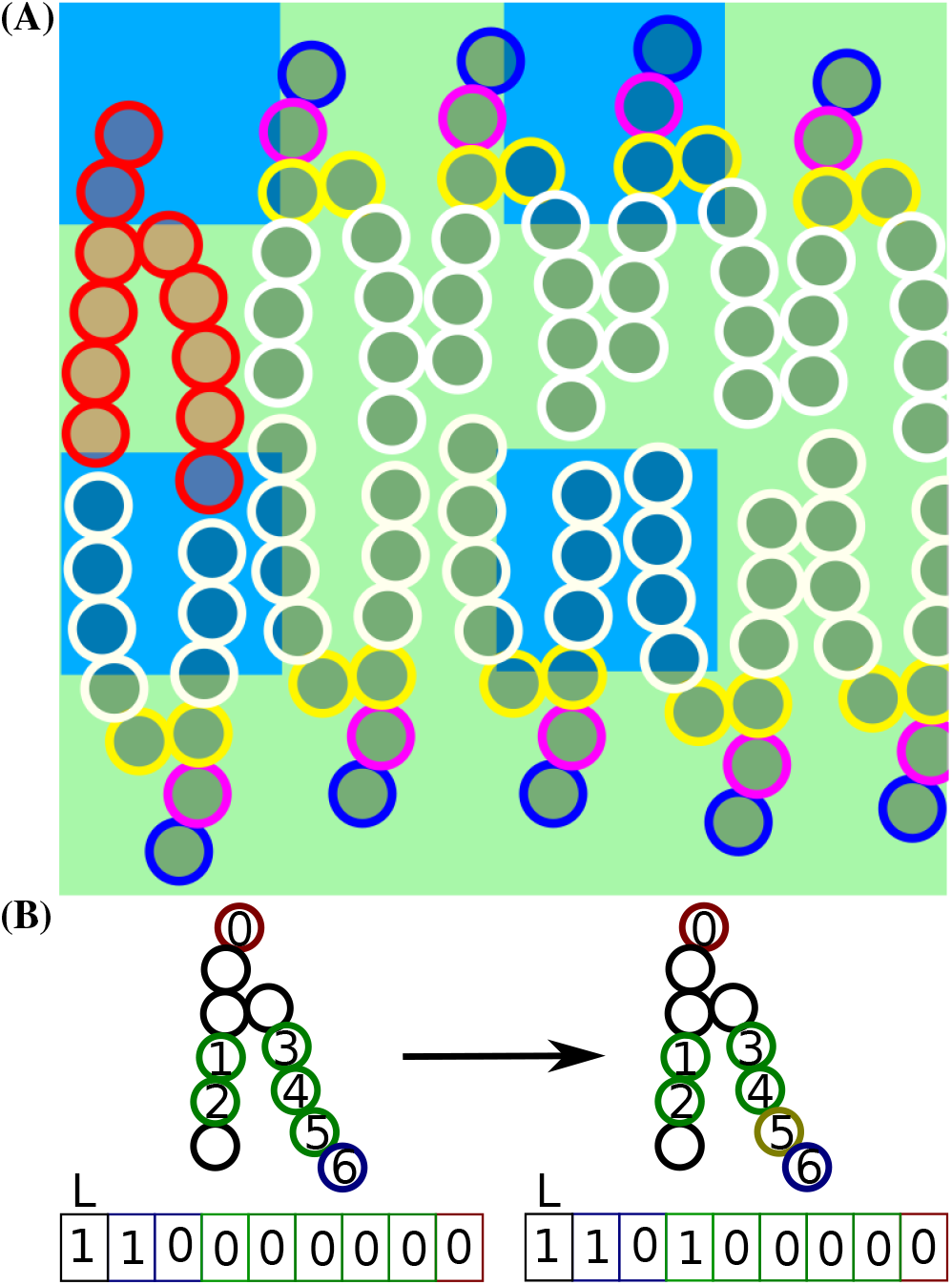
SGC algorithm employed. A) 2D representation of the checkerboard decomposition for a lipid membrane; lipids are drawn in MARTINI representation with standard MARTINI coloring, checkboard is in green and active cells in blue. A random particle is chosen within each active cell for an alchemical transformation. Since the red molecule stretches across multiple active cells, it is pathological and can lead to data races. B) Calculation of the molecule hash for a typical molecule in a bilayer, with colour indicative of SGC repre-sentation. Every bead in the molecule is assigned an offset in the hash so that changes in hash can be directly computed by changing the relevant bits. For instance, an alchemical transformation of bead labeled 5 from green (state 0) to gold (state 1) results in a change of bit 5 of the hash. Discrete, finite-valued collective variables such as leaflet side can be directly incorporated into the hash as well, represented here with a value of 1 for the upper leaflet.

To associate a chemical potential to a given molecule, we need to assign a unique number—a hash—to a given chemical structure. This hash needs to include collective variables, such as leaflet, if they are relevant to chemical potential values. To construct this hash, we simply aggregate all potential chemical states of beads in a molecule. As a relevant example, let us consider the coarse-grained lipid depicted in Fig. 1B. For this particular lipid, which we depict using the coarse-grained MARTINI force field (4), seven beads can change their chemical type. First the headgroup (red) can change between PC (Q_0_ MARTINI beadtype) and PE (Q_d_). The green beads can either correspond to saturated (C_1_) or unsaturated (C_3_) states. The last bead, in blue, is used to change the length of the acyl tail: it can either be saturated, unsaturated or “ghost” (empty). The hash is constructed from the minimal binary representation: a two-state bead occupies one bit, while a three-state bead occupies two bits. Some of the hash values may correspond to unphysical states, for instance the same binary representation is used for both three- and four-state beads. Additionally, some states may be chemically unavailable, e.g. non-contiguous unsaturations in biologically relevant lipids. Both unphysical and chemically unavailable states are assigned a chemical potential *μ* = −∞ to forbid anyalchemical transformation involving these states.

The Monte-Carlo procedure (see Alg. 1) is similar to (11), but require a few more memory transactions. Effectively, after picking a random particle, the algorithm must resolve to which molecule the particle belongs, followed by retrieving the hash of the molecule. A new random state for the particle is then generated, as well as its associated hash. Computing the new hash requires resolving the hash offset of the current bead. Additionally, this procedure implies that parallel transformations on the same molecule result in data races for hashes — read and write commands occurring at the same time from multiple threads resulting in corrupt data states. Therefore, large molecules, which span multiple active cells (see red molecule on Fig 1), are pathological. To solve this, we add a molecular lock to prevent multiple changes to the same molecule within a single Monte-Carlo step.

### Algorithm 1 Monte-Carlo Procedure

~~~
**for** c in active cells **do**
     *p* ← random particle ∈ *c*
     mol ← MoleculeIndex[*p*]
     molHash ← Hash [mol]
     *o* ← Offset [*p*]
     *s* ← States [*p*]
     *s′* ← Random state ≠ *s*
     molHash’ ← (Hash & ~ (mask ≪ *o*))|(*s′* ≪ *o*)
     Δ*U* ← *U*[*s′*] – *U*[s]
     Δ*μ* ← *μ* [molHash’] – *μ*[molHash]
     **if** *R*(0, 1) < exp(-*β*(Δ*U* – Δ*μ*)) **then**
          lock ← atomicCAS(&Lock[mol], 0, 1)
          **if** lock **then return**
          Hash [mol] ← molHash’
          States [μ] ← *s′*
~~~

The base-2 representation for chemical states ensures high numerical performances as alchemical changes can be directly computed through bitwise operations. This comes at the cost of chemical space; for instance, in Fig 1B, the fourth state of the blue bits (11) does not represent a meaningful physical state. Since we use 32-bit integers for hashes, the worst case scenarios involve either losing a chemical state every two bits (e.g. a molecule composed of only blue bits in Fig1), leading to 3^16^ = 4.3 · 10^6^ chemical species, or using beads with more than 2^16^ + 1 = 65537 chemical states; in which case there can be only a single such bead per molecule. To our knowledge, no simulation has attempted mixtures of more than 2^16^ components yet and we believe that this is sufficient.

If the chemical potential is dependent on a collective variable, for instance if chemical potentials are different on the two leaflets of a membrane, then the system is out of equilibrium. These systems can exhibit peculiar properties, such as net flows, which tend to depend on kinetic rates in the system. In order to use chemical potentials to describe the system, the natural relaxation timescale of the system (e.g. flip-flop for asymmetric phospholipids bilayers) must be much longer than simulation timescales. If the natural relaxation timescale is similar to simulation timescales, then the system will exhibit properties that are dependent on simulation condition choices, and particularly the SGC relaxation timescale which creates the out-of-equilibrium conditions.

## RESULTS

In order to demonstrate the value of our method, we take a look at a biologically relevant system: a membrane with an asymmetric lipid composition. We simulate a membrane comprised of PC and PE. On the lower leaflet, we impose Δ*μ* = 0 between any two chemical species. This results in a higher proportion of PE present due to hydrogen bonds forming between their headgroups, with ≈ 88% of lipids being PE. On the upper leaflet, we impose a difference between PC and PE molecules of Δ*μ* to proxy effects of P4-ATPase proteins. As outlined in the introduction, this assumes that P4-ATPA_SE_ binds all PC molecules equally, independently of acyl tail nature.

To measure asymmetry, we define the headgroup asymmetry parameter 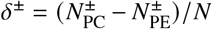, which measures how different the headgroup populations are on each leaflet (see Fig2). As expected *δ*^+^ shows a sigmoid-like behavior, where the free energy is largely dominated by the mixing entropy at large values of Δ*μ*. The value of *δ*^±^ = 0 is not reached at Δ*μ* = 0, due to hydrogen bonding occurring between PE heads. The composition of the lower leaflet barely changes, indicative of absence of coupling between headgroup compositions of both leaflets. The two curves intersect at Δ*μ* = 0, as expected.

Beyond resulting in asymmetric membranes, this simulation also yields an important result: P4-ATPA_SE_ proteins need to exert 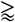 20 kJ/mol of work on lipids to create a strongly PC-dominated upper leaflet. This value is compatible with free energy release during hydrolysis of ATP (≈ 30 kJ/mol).

This algorithm has 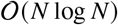 time-complexity since the amount of Monte-Carlo attempts in a single step grow linearly with system size. This means that it can be used to study large-scale systems, for instance to do finite-size scaling of critical membranes. Additionally, if the membrane has only a single SGC ensemble and no unregulated components — molecules whose chemistry cannot change, e.g. cholesterol —, equilibration becomes independent of long-range diffusion. This is similar to the molecular dynamics coupled with alchemical steps, where two lipid chemical states are swapped in composition-conserving non-equilibrium transformations (15, 16). However, only a single non-equilibrium move can be attempted per update, which in turn implies a 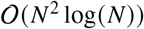 time complexity.

**Figure 2:**
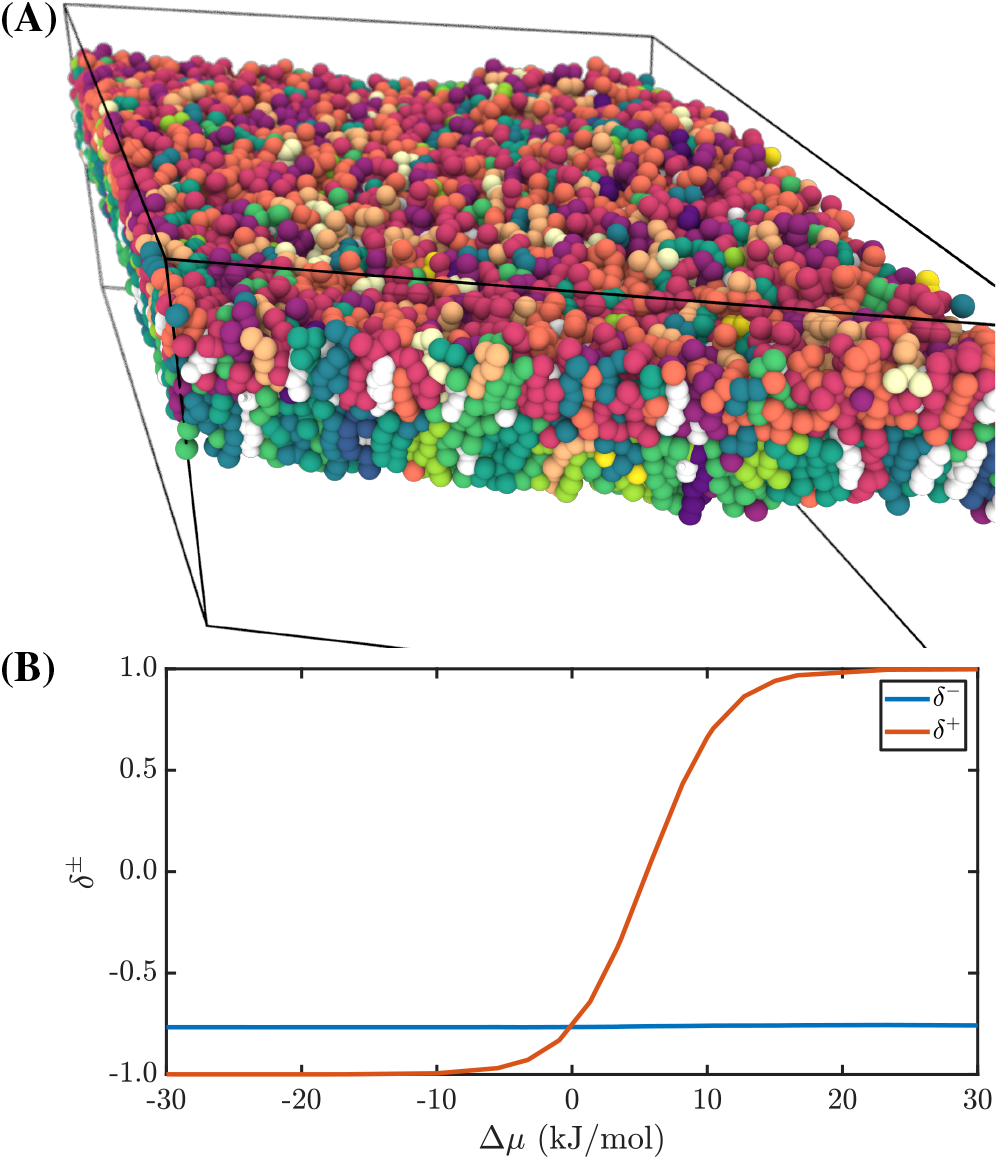
Asymmetric membrane properties. A) Snapshot of a typical configuration at Δ*μ* = 10 kJ/mol. Cholesterol is coloured in white, while PC and PE are coloured according to their unsaturation level on different color scales to differentiate them. B) Resulting headgroup asymmetry *δ*^±^. At Δ*μ* = 0, PE molecules dominate both layers, with ≈ 88% of molecules being PE. The composition of the lower leaflet is nearly unaffected by the changes of the upper leaflet.

## CONCLUSION

We developed a molecule-wise SGC algorithm that enables simulation of lipid membranes with distinct sets of chemical potentials on different leaflets. This results in membranes with asymmetric composition between the two leaflets. We hope that the simulation tools deployed here will enable research into regulated ensembles proposed in (9) and into properties of asymmetric membranes.

## AUTHOR CONTRIBUTIONS

M.G. and T.B. designed the research. M.G. wrote the software, carried out simulations, analyzed the data. M.G. and T.B. wrote the article.

## ACKNOWLEDGMENTS

We thank Nikita Tretyakov for a critical reading of this manuscript. This project was supported by the Alexander von Humboldt-Stiftung (AvH) and the Deutsche Forschungsgemeinschaft (DFG). We acknowledge usage of computational resources from the Max-Planck Computing and Data Facilities (MPCDF).

## SOFTWARE

Molecular dynamics simulations make use of the HOOMD-Blue engine (13, 14, 17), a DPD thermostat (18) and the MARTINI force-field (4). Initial topologies are built using the hoobas molecular builder (19). The SGC HOOMD-Blue plugin for HOOMD-version 2.9.3 is available in supplementary material, as well as on https://gitlab.mpcdf.mpg.de/mgirard/SGC-molecules.

## SUPPLEMENTARY MATERIAL

An online supplement to this article can be found by visiting BJ Online at http://www.biophysj.org.

